# Noisy Perturbation Models Distinguish Network Specific from Embedding Variability

**DOI:** 10.1101/550467

**Authors:** A. Piehler

## Abstract

Recently, measurement technologies allowing to determine the abundance of tens signaling proteins in thousands of single cells became available. The interpretation of this high dimensional end-point time course data is often difficult, because sources of cell-to-cell abundance variation in measured species are hard to determine. Here I present an analytic tool to tackle this problem. By using a recently developed chemical signal generator to manipulate input noise of biochemical networks, measurement of state variables and modeling of input noise propagation, pathway-specific variability can be distinguished from environmental variability caused by network embedding. By employing different sources of natural input noise, changes in the output variability of the apoptosis pathway were measured, indicating that also synthetic noisy perturbations are biologically feasible. The presented analytic tool shows how signal generators can improve our understanding of the origin of cellular variability and help to interpret multiplexed single cell information.

## Introduction

Chemical reactions occurring in living cells often include small molecule numbers. The birth-death processes of these molecules can be accurately described by the Chemical Master Equation (CME) [1, 2]. The Fokker-Planck equation is an approximation of the master equation and can be solved for large systems of reactions such as signaling pathways [3, 4, 5]. One complication in comparing models of such pathways with experimental data is that not all species inside the cell can be measured simultaneously [6, 7]. Many processes representing the embedding are thus lumped into rate constants and not modeled explicitly. In reality rate constants vary from cell to cell and over time and these fluctuations will contribute to the observed variability in the state variables of a model [8].

To solve this problem, Elowitz *et al.* devised the two color method that allows to dissect variability of rate constants from fluctuations of the state variables [9, 6, 10]. The two color method exploits that two copies of the same molecular circuit will experience the same fluctuations in rate constants and can thus be distinguished from uncorrelated noise in the state variables[11]. Since placing a second copy of a circuit into a cell is only possible for very simple circuits, the method is inherently limited.

On the other hand, newly developed measurement technologies such as CyTOF, CyCIF, and 4i provide unprecedented experimental power and make it possible to routinely measure more than 40 species in single cells, allowing to monitor larger reaction networks [12, 13, 14]. Multiplexed RNA profiling even allows to monitor hundreds of transcripts in the same single cell [15]. The drawback of these methods is that they only allow endpoint measurements. Whereas live-cell measurements observe the same cell and thus the same embedding over time, endpoint time-course data consists of different cells measured at each time point, implying that network embeddings are also different. Currently, it is hard to quantify the source of variability in end-point data-sets, because methods such as the dual reporter system are missing.

Recent advances in microfluidic technologies make it possible to generate and reproduce stochastic input trajectories *I*(*t*) of a chemical signal with precisely controlled variability and autocorrelation [16]. Stochastic dynamic signals can be used to probe network properties of biological systems (see Figure 1) such as dynamic correlations [17, 18, 19], noise propagation [20, 21], filtering properties of network components [22] and stochastic resonance (personal communication Savas Tay). Stochastic perturbations of particular nodes in the network by e.g. small molecules can also enhance identifyability of network topologies and rate constants [8, 23, 24, 25, 26, 27].

**Figure 1:**
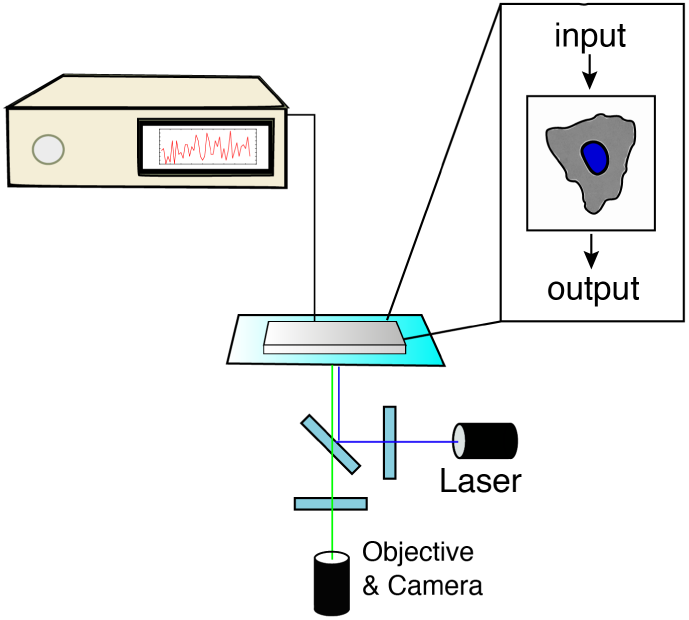
Experimental setup for input control and output measurement: Cells cultured in a microfluidic device can be stimulated dynamically with a noisy input signal provided by the universal chemical signal generator [16]. The cellular response can then be measured by a fluorescence microscope that detects levels of fluorescent molecules such as GFP and Alexafluor.

Here I introduce a concept, that allows to dissect pathway specific from embedding variability by using a chemical signal generator and measurement of output molecule levels. The method is introduced using a model with one input variable *I*, one embedding variable *E* and one measured output *X*. The principle extends to larger systems with many state-variables and rate constants, as it does not require a circuit copy. By accounting for embedding variability rigorous comparison of stochastic models with network specific variability becomes also possible for end-point time course measurement data.

In the toy model considered in detail below, one can think of the generated input signal as a receptor ligand and the output active receptor levels *X*, where the embedding consists of other factors, that contribute to receptor activation or deactivation, that one does not model explicitly (e.g. Kinases, Cofactors, etc.) and are lumped into rate constants (see Figure 2).

**Figure 2:**
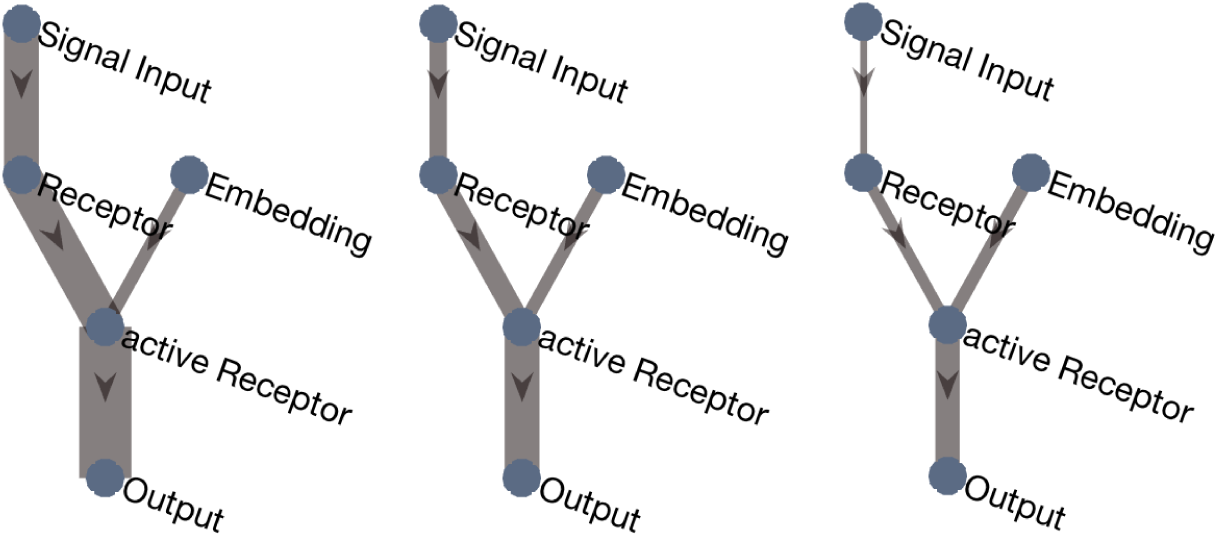
The signal generator can produce stochastic traces with controlled variability of the ligand *I* and allows to manipulate the input noise amplitude and autocorrelation. The input noise will propagate along the pathway according to biochemical parameters and leave other incoming sources of variability unchanged. This allows to distinguish pathway specific from embedding variability.

## Modeling noisy stimulation

Let us study a toy model where the production rate of a species *X* depends on the input *I* and some embedding factor *E*.

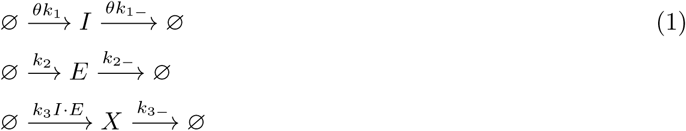

If one can control the rates of the input signal *I* it is possible to quantify the contribution of an external factor *E* to the total variability in the output *X* by measuring *X* only. To control the timescale and hence the variability in *I*, without affecting its mean level, we introduce the input turnover parameter *θ*, in both the production and degradation rate. The stochastic trajectory for *I* can be simulated with a stochastic simulation algorithm (e.g. the Gillespie algorithm) and reproduced by the signal generator.

## CME of the reaction system

For the considered system 1 the CME reads

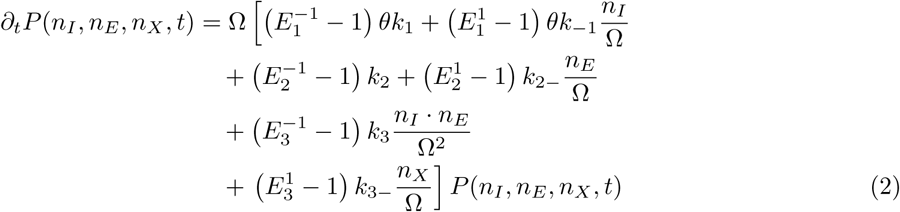

where *P* (*n_I_, n_E_, n_X_, t*) is the probability density, Ω the system size and *E^−S_ij_^* the step operators. By using Van Kampen’s method one can derive a Fokker-Planck equation for 2 (see APPENDIX). Multiplying the Fokker-Planck equation by the stochastic variables *ϵ_i_ϵ_j_* and integrating over the whole domain, one gets a Matrix Differential Equation for the covariance time evolution 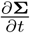. In steady state 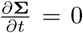, the resulting Lyapunov Equation (see APPENDIX) for the covariance can be readily solved with a computer algebra software. The solution for the noise is given by the covariance normalized by the deterministic concentrations *〈φ_i_〉〈φ_j_〉*. We get for the noise in the output *X*

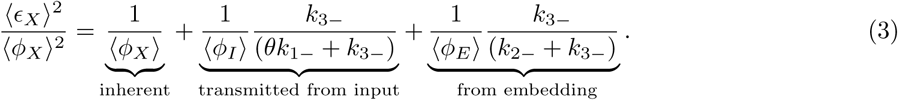

Analyzing 3, one can appreciate, that including the parameter *θ* in both the production and degradation rate allows to change the timescale of the input reaction and hence the transmitted noise, without affecting the mean concentrations. The faster the rates and hence the fluctuations of the input noise compared to the fluctuations of the output, the more noise will be filtered (see Figure 3). For very fast fluctuations the transmitted input noise converges to zero.

**Figure 3:**
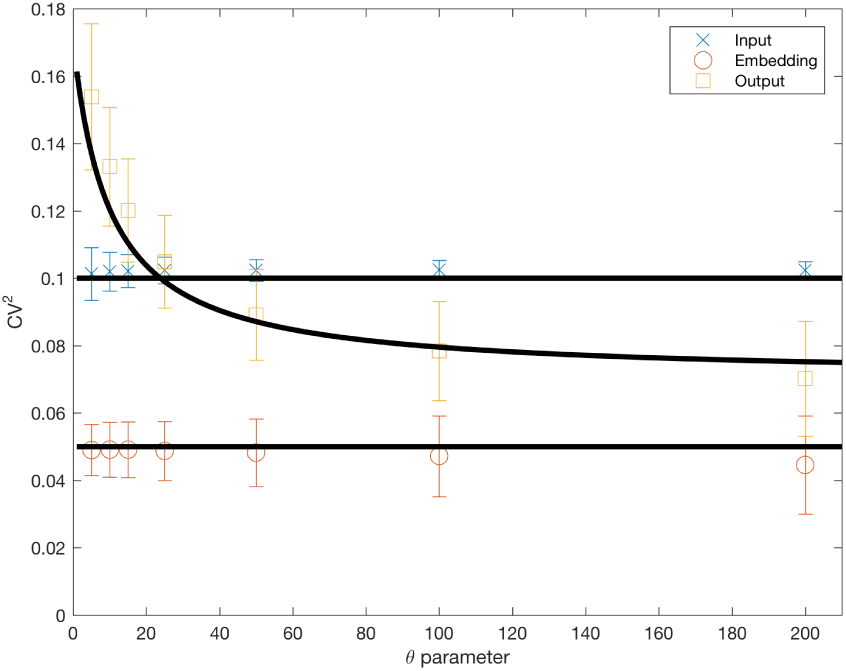
Black: Linear Noise Approximation; Color: Gillespie Simulation; Parameters: Ω = 1*, k*_1_ = 10^*−*5^*, k*_1*−*_ = 10^*−*6^*, k*_2_ = 2 * 10^*−*5^*, k*_2*−*_ = 10^*−*6^*, k*_3_ = 2 * 10^*−*6^*, k*_3*−*_ = 10^*−*5^; Error bars indicate variability between simulation runs. The input noise that is propagated to the output depends on the timescale difference at which reactions occur. The timescale of the input can be changed by the parameter *θ*. With increasing turnover *θ* more and more noise is filtered. In the limit *θ* → ∞ the contribution of input noise to the measured variability in the output vanishes.

On the other hand changing only the production rate and hence the mean levels of *I*, can be used to tune the amplitude of the input noise. The higher the mean levels 〈*n_I_*〉, the lower gets the input noise and hence the noise transmitted to the output (see Figure 4).

**Figure 4:**
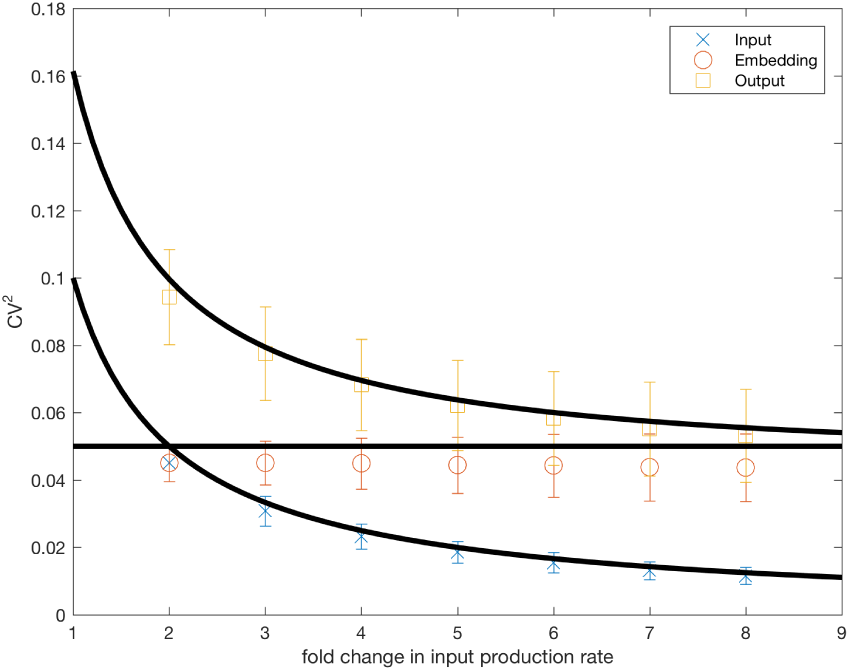
Black: Linear Noise Approximation; Color: Gillespie Simulation; Parameters: Ω = 1*, k*_1_ = 10^*−*5^*, k*_1*−*_ = 10^*−*6^*, k*_2_ = 2 * 10^*−*5^*, k*_2*−*_ = 10^*−*6^*, k*_3_ = 2 10^*−*6^*, k*_3*−*_ = 10^*−*5^; Error bars indicate variability between simulation runs. The input noise that is propagated to the output depends on the amplitude of the input noise. With increasing levels of *I* the amplitude and hence the transmitted noise is decreasing. In the limit *I → ∞* the contribution of input noise to the measured variability in the output vanishes.

While the network specific variability contribution 4

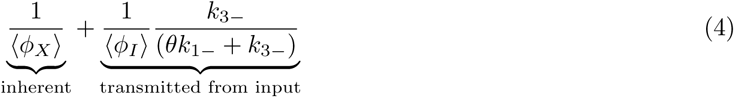

to the total variability in the output *X* is changing with the input characteristics, the embedding variability contribution

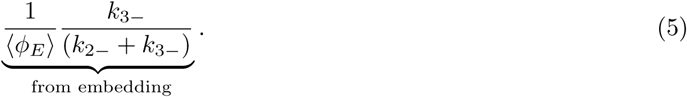

stays unaffected. Generating input noise with changing, but known turnover rates *θ* or input noise amplitudes 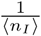, one can measure the changes in the output variability. Measurement of the absolute number of molecules of the output variable *X* allows to calculate the inherent contribution 1*/〈n_x_*〉 to the network specific noise. The transmitted input noise can be determined by fitting the second term of 4 to the data. Finally the embedding variability can be determined as the constant contribution after subtracting the network specific variability from the total output variability.

This dissection of network specific and embedding variability is essential in model selection problems, where embedding variability is otherwise confounding the network inference process.

## Autocorrelation of the system output

Temporal correlations in the output signal can be used to indentify sources of noise [28, 7]. Here this notion and the signal generator as a noise-source are used to quantify embedding autocorrelation in live-cell data. The autocorrelation 〈*ϵ_i_*(*t*)*ϵ_i_*(*t* + *τ*)〉 of a system is a measure of how the variable *ϵ_i_* at time *t* correlates with itself at a later time point *t* + *τ* and hence depends on the timescale at which fluctuations occur. For the normalized autocorrelation in the output variable *ϵ*_3_ of reaction system 1, one finds equation 6

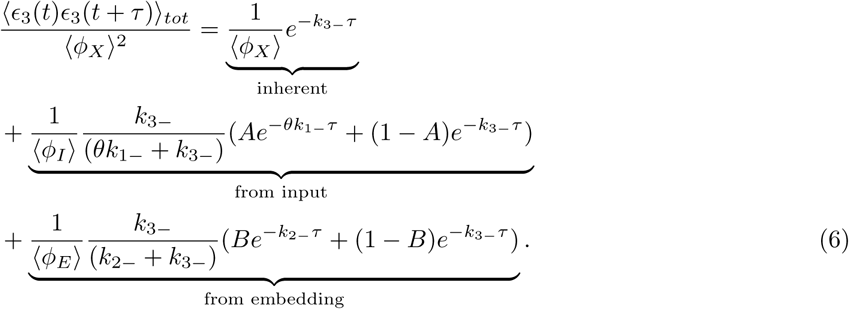

where integration constants have been chosen, such that for *τ* = 0 the normalized autocorrelation matches the normalized covariances. In this case *A* and *B* result to 7 and 8, respectively (see also Supplementary Information for details).

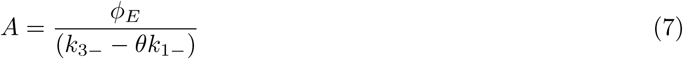

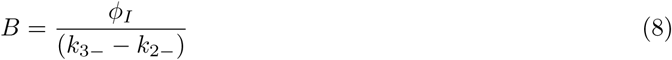

The solution 6 can be interpreted the following. The term 9 is network specific autocorrelation

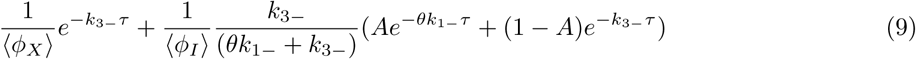

and its first term is similar to the autocorrelation of a simple production and degradation processes, such as reactions for *I* and *E* and can thus be interpreted as inherent autocorrelation. The second term of 9 stems from auto-cross-correlations involving 〈*ϵ*_1_(*t*)*ϵ*_3_(*t* + *τ*)〉 (see Supplementary) and represents autocorrelation transmitted from the input.

The embedding autocorrelation 10 corresponds to the third term in equation 6

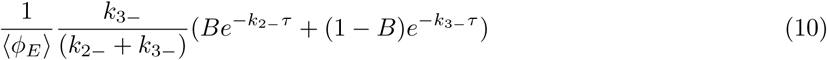

originates from the auto-cross-correlation 〈*ϵ*_2_(*t*)*ϵ*_3_(*t* + *τ*)〉 and is transmitted from the embedding.

## Apoptosis Network Embedding

To test if synthetic noisy stimulation can be biologically feasible and to illustrate the network embedding principal, variability in cell death caused by two distinct pathways was studied. HeLa cells undergo apoptosis in response to both intrinsic and extrinsic death signals. In response to both signals, the molecule BID is cleaved to truncated BID (tBID), which leads to BAK and BAX pore formation in the outer mitochondrial membrane in a process known as Mitochondrial Outer Membrane Permeation (MOMP), resulting in cell death [29]. The apoptosis network receives inputs from the intrinsic death pathway by PIDD (p53-induced death domain protein), which leads to BID truncation via Caspase-2 upon genotoxic stress [30, 31]. The extrinsic death pathway gets activated by TNF receptor stimulation, which leads to receptor internalisation and truncation of BID by Caspase-8 [32, 33]. HeLa cells were stimulated with TRAIL to activate extrinsic apoptosis and cisplatin to activate intrinsic apoptosis. Although stimulation is constant, the signals received by BID are shaped by the intrinsic and extrinsic death pathway and thus stochastic. A live cell movie of hundreds of stimulated cells was recorded, nuclei were tracked throughout time and the time of death determined for each single cell. For both treatments a large cell-to-cell variability in the time of death can be observed (see Figure 5).

**Figure 5:**
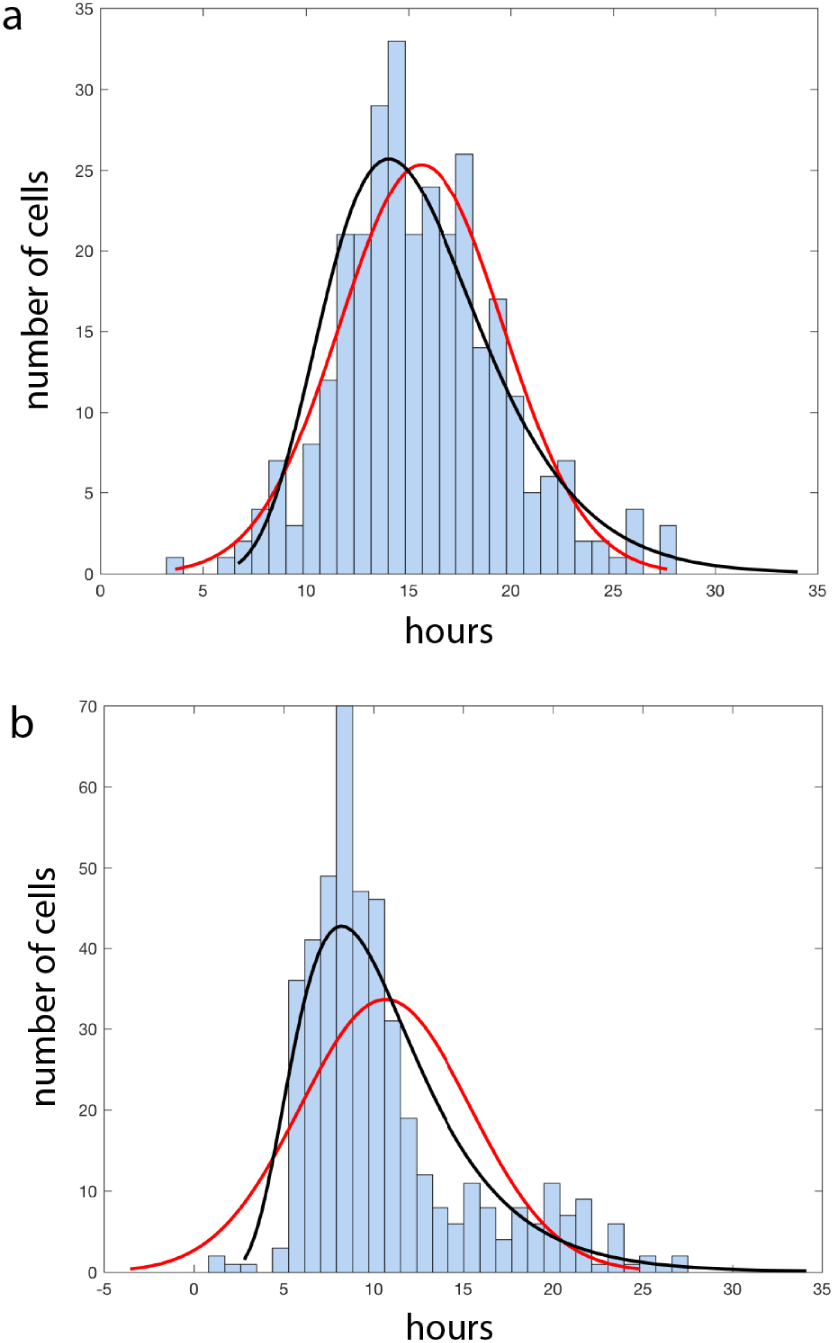
Histograms for the time of death post stimulation with a) 1*µg* TRAIL and b) 60*µg* cisplatin. Gaussian (red) and log normal distributions (black) were fitted to the histograms. The mean time of death for cisplatin is 10,7h *±* 4,7h (noise 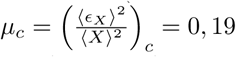 and for TRAIL 15,6h *±* 4h (noise 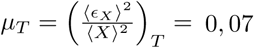 post-stimulation. The TRAIL histogram is normally distributed and the cisplatin histogram shows a tail towards high death times.

In this example, the BID node receives inputs from two natural, but distinct noisy signals, the intrinsic and extrinsic death pathway, respectively. While the contribution of the embedding (e.g. variability in BAC and BAX) to the variability in death times is constant, the network specific variability depends on the BID input variability. The difference in the input noise of these pathways can be observed at the output (*µ_c_* = 0, 19 vs. *µ_T_* = 0, 07), which in this example is the absolute time of cell death (Figure 5). This shows that natural input signal variability is faithfully transmitted to the output and supports that also synthetic stochastic signals lead to measurable variability differences in the output. While the characteristics of natural input noise sources are unknown, synthetic signals are known precisely, can be tuned to generate data-sets optimal to fit curves such as shown in figure 3 as well as 4 and can be easily adapted for new targets. Future microfluidic devices, that allow noisy perturbations and multiplexed state measurements, together with analytic tools as presented here, promise a powerful approach to improve cell fate engineering.

## Discussion

In this work the need for an analytic noise dissection tool for large data-sets arising form single-cell end-point measurements has been addressed. The presented method to dissect noise of larger pathways is based on input noise propagation along the pathway that leaves other incoming sources of variability that merge with the pathway unchanged.

Rate constants reflect the environment in which a circuit is embedded. As such the observed embedding variability in these rates can be caused by deterministic or stochastic sources [34]. For example, the population context (cell density and morphology) and nonlinear reactions, such as the the cell cycle are deterministic sources of variability [35, 36, 37, 38], while variability in many protein levels is of stochastic nature [32, 39, 40, 41].

Network Embedding Analysis is a powerful tool to find hidden deterministic variables causing embedding variability. Nodes, where the largest embedding variability is observed might point to a regulatory mechanism. By further investigating these nodes, one can start to systematically identify the underlying regulatory mechanisms. Thus, the presented method provides means, by which cellular information processing can be understood and the fraction of cells reprogrammed to a particular fate, can be systematically improved.

Reaction mechanisms and topologies of embedded networks are often unknown. The presented reaction system is one model used here to illustrate the power the signal generator in quantifying and dissecting noise. In order to understand the topology and dynamics of networks, models needs to be inferred from the data[42]. This process will be more reliable, if the fitted model is not confounded by embedding variability and thus greatly enhanced by employing the signal generator.

The cellular phenotype is intimately linked to biochemical network topology and rate parameters. Ultimately a quantitative understanding of these networks is necessary, if we want to reliably engineer single cell fate.

## Methods

### Cell Culture and Drug Treatment

HeLa cells were cultured for two days in Dulbecco’s Modified Eagle’s Medium (Sigma Aldrich) with 10% Fetal Bovine Serum (Sigma Aldrich) at 37C and 4% CO_2_ before seeding in a 96 well plate (Greiner). Cells were then cultured over night, thereafter stimulated with 60*µ*g cisplatin or 1*µ*g TRAIL and imaged over 25 hours.

### Imaging and Cell Tracking

A Nikon Ti Eclipse Microscope has been used for automated fluorescence time lapse microscopy. Cells were stained with the nuclear dye Hchst (ThermoFisher), imaged every 40min and subsequently tracked with CellProfiler. Cell death was indicated by a sudden switch-like increase in Hchst signal. To extract the time of death of hundreds of cells reliably in automated manner, hill curves were fitted to each Hchst time course and the time of death parameter extracted (see Supplementary Information).

### Gillespie Simulation

Gillespie Simulations have been performed with a custom Matlab algorithm using the direct method and verified by comparison to Nezar Abdennur’s Github algorithm “Gillespie Stochastic Simulation Algorithm” with comparable results. For details on the simulation parameters and time course results, see the Supplementary Information.

## Appendix

### Step operators

Application of step operators *E^−S_ij_^* increase the *i*-th species of the *j*-th reaction by the stoichiometric coefficient *S_ij_*. Its Taylor expansion to order *O*(Ω^*−*3*/*2^) is

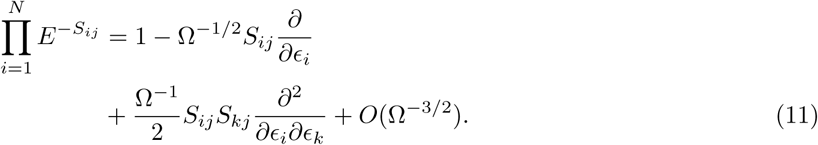

## Derivation of the Covariances

The Fokker-Planck equation derived from 2 is

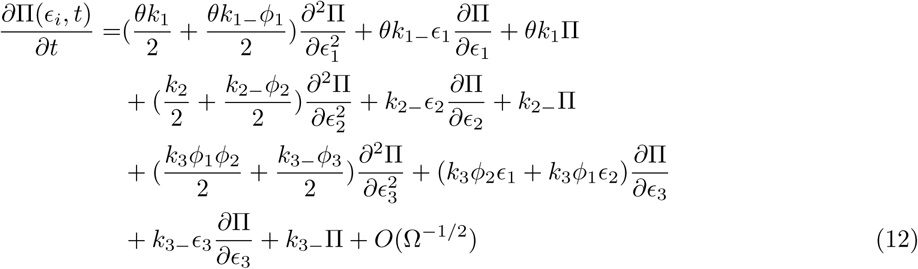

(see SI for a detailed derivation). From this Fokker-Planck equation 12, the second moments can be calculated by multiplying equation 12 by the *∊_i_∊_j_* and integrating over the whole domain to get the Matrix Differential Equation

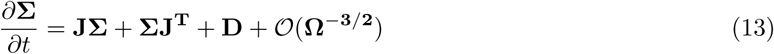

In steady state 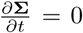, equation 13 becomes a Lyapunov equation and can be solved for the covariances.

## Acknowledgment

I thank Ramon Grima, Peter Swain and Christoph Zechner for feedback on the manuscript and acknowledge Lucas Pelkmans for providing experimental resources.

