## Supplementary for "Noisy Perturbation Models Distinguish Network Specific from Embedding Variability"

A. Piehler[a,1]

[a]Interdisciplinary Systems Biology PhD Program ETH Zurich and University of Zurich, 8057 Zurich, Switzerland
[1]Institute for Molecular Life Sciences, 8006 Zurich, Switzerland

February 14, 2019

### A noisy stimulation process

Let us consider the following reaction system

$$\begin{aligned}
&\varnothing \xrightarrow{\theta k_1} I \xrightarrow{\theta k_{1-}} \varnothing \qquad (1)\\
&\varnothing \xrightarrow{k_2} E \xrightarrow{k_{2-}} \varnothing \\
&\varnothing \xrightarrow{k_3 I \cdot E} X \xrightarrow{k_{3-}} \varnothing
\end{aligned}$$

where the production of $X$ depends on both the input $I$ and some embedding factor $E$.

### Deterministic description of the system

The deterministic rate equations describing this system are given by

$$\begin{aligned}
\frac{d\phi_I}{dt} &= \theta k_1 - \theta k_{1-}\phi_I \qquad (2)\\
\frac{d\phi_E}{dt} &= k_2 - k_{2-}\phi_E \\
\frac{d\phi_X}{dt} &= k_3 \cdot \phi_I \cdot \phi_E - k_{3-}\phi_X
\end{aligned}$$

where $\phi$ denotes the mean concentrations. In steady state the concentrations are $\phi_I = \frac{k_1}{k_{1-}}$, $\phi_E = \frac{k_2}{k_{2-}}$ and $\phi_X = \frac{k_1 \cdot k_2 \cdot k_3}{k_{1-} k_{2-} k_{3-}}$. As desired the mean concentration $\phi_I = \frac{k_1}{k_{1-}}$ is not influenced by $\theta$.

### Chemical Master Equation

In general one can write a system of $R$ chemical reactions as

$$s_{1j}X_1 + ... + s_{Nj}X_N \xrightarrow{k_j} r_{1j}X_1 + ... + r_{Nj}X_N \qquad (3)$$

where $X_i$ denotes the $i$-th chemical species and $j$ is the reaction index running from 1 to $R$. $s_{1j}$ and $r_{1j}$ denote the stoichiometric coefficients and $k_j$ is the macroscopic reaction rate. The vector

$\vec{n}$ contains the absolute number of molecules of each species. One can write the Chemical Master Equation (CME) [1] associated with system 1 as:

$$\partial_t P(\vec{n},t) = \Omega \sum_{j=1}^{R} \prod_{i=1}^{N} \left(E^{-S_{ij}} - 1\right) \hat{f}_j(\vec{n},\Omega) P(\vec{n},t) \tag{4}$$

$\Omega$ denotes the volume or in other words the size of the system. For the bespoken reaction system 1 the stoichiometric matrix is given by the $3 \times 6$-matrix. The stoichiometric matrix is defined by $S_{ij} = r_{ij} - s_{ij}$ [2].

$$\mathbf{S} = \begin{bmatrix} 1 & -1 & 0 & 0 & 0 & 0 \\ 0 & 0 & 1 & -1 & 0 & 0 \\ 0 & 0 & 0 & 0 & 1 & -1 \end{bmatrix} \tag{5}$$

The function $f_j(\vec{n},t)$ is known as propensity and depends on the molecule numbers $n_i$ as well as the stoichiometric coefficients $s_{ij}$ and the volume $\Omega$. According to the law of mass action

$$\hat{f}_j(\vec{n},\Omega) = k_j \prod_{z=1}^{N} \Omega^{-s_{zj}} \frac{n_z!}{(n_z - s_{zj})!}. \tag{6}$$

For the studied reaction system 1 the propensities are $f_1 = \theta k_1$, $f_2 = \theta k_{1-} \frac{n_1}{\Omega}$,$f_3 = k_2$, $f_4 = k_{2-} \frac{n_2}{\Omega}$,$f_5 = k_3 \frac{n_1 n_2}{\Omega^2}$, $f_6 = k_{3-} \frac{n_3}{\Omega}$. Further, $E^{-S_{ij}}$ is known as the step operator, which can be Taylor expanded in powers of the square root of the system size

$$\prod_{i=1}^{N} E^{-S_{ij}} - 1 = -\Omega^{-1/2} S_{ij} \frac{\partial}{\partial \epsilon_i} + \frac{\Omega^{-1}}{2} S_{ij} S_{kj} \frac{\partial^2}{\partial \epsilon_i \partial \epsilon_k} + O(\Omega^{-3/2}). \tag{7}$$

### CME of the reaction system

For the considered system 1 the CME reads

$$\begin{aligned} \partial_t P(n_1,n_2,n_3,t) = \Omega \Big[ & \left(E_1^{-1} - 1\right) \theta k_1 + \left(E_1^{1} - 1\right) \theta k_{-1} \frac{n_1}{\Omega} \\ & + \left(E_2^{-1} - 1\right) k_2 + \left(E_2^{1} - 1\right) k_{2-} \frac{n_2}{\Omega} \\ & + \left(E_3^{-1} - 1\right) k_3 \frac{n_1 \cdot n_2}{\Omega^2} + \left(E_3^{1} - 1\right) \frac{k_{3-} n_3}{\Omega} \Big] P(n_1,n_2,n_3,t) \end{aligned} \tag{8}$$

### Derivation of the LNA from the CME

An approximate solution to the CME 8 can be found by using Van Kampen's method, which leads to the linear noise approximation (LNA) of the Master Equation. To this end, one can plug the expanded step operator 7 and the propensities 6 into the CME 4 after transforming the latter into continuous variables. The change of the discrete variables $n_i$ into continuous variables $\epsilon_i$ is achieved by using Van Kampen's ansatz [1]

$$\frac{n_i}{\Omega} = \phi_i + \Omega^{-1/2} \epsilon_i, \tag{9}$$

where $\phi$ is the deterministic concentration and $\epsilon$ the stochastic variable.

$$P(n_1,n_2,n_3,t) = P(\Omega\phi_1 + \Omega^{1/2}\epsilon_1, \Omega\phi_2 + \Omega^{1/2}\epsilon_2, \Omega\phi_3 + \Omega^{1/2}\epsilon_3, t) = \Omega^{1/2}\Pi(\epsilon_1,\epsilon_2,\epsilon_3,t) \tag{10}$$

Using the time derivative of 9 $\partial_t \epsilon = -\Omega^{1/2}\partial_t \phi$, the transformation from discrete molecule numbers to the continuous stochastic variables is given by

$$\frac{\partial}{\partial t}P(n_1,n_2,n_3,t) = \frac{\partial \Pi(\epsilon_i,t)}{\partial t} - \sum_i \Omega^{1/2}\frac{\partial \Pi(\epsilon_i,t)}{\partial \epsilon_i}\frac{\partial \phi_i}{\partial t}. \tag{11}$$

We can explicitly write down the step operators for the reaction system of interest 1 by plugging in stoichiometric coefficients into 7. For each species $i$ we get

$$\begin{aligned}(E_i^1 - 1) &= \Omega^{-1/2}\frac{\partial}{\partial \epsilon_i} + \frac{\Omega^{-1}}{2}\frac{\partial^2}{\partial \epsilon_i^2} + O(\Omega^{-3/2}) \\ (E_i^{-1} - 1) &= -\Omega^{-1/2}\frac{\partial}{\partial \epsilon_i} + \frac{\Omega^{-1}}{2}\frac{\partial^2}{\partial \epsilon_i^2} + O(\Omega^{-3/2}).\end{aligned} \tag{12}$$

Plugging in the propensities 6, the ansatz 9 and the expanded step operators 12 into 11, we get the following equation

$$\begin{aligned}\frac{\partial}{\partial t}P(n_1,n_2,n_3,t) &= \frac{\partial \Pi(\epsilon_i,t)}{\partial t} - \sum_i \Omega^{1/2}\frac{\partial \Pi(\epsilon_i,t)}{\partial \epsilon_i}\frac{\partial \phi_i}{\partial t} \\ &= \Omega\Bigg[\left(-\Omega^{-1/2}\frac{\partial}{\partial \epsilon_i} + \frac{\Omega^{-1}}{2}\frac{\partial^2}{\partial \epsilon_i^2}\right)\theta k_1 + \left(\Omega^{-1/2}\frac{\partial}{\partial \epsilon_1} + \frac{\Omega^{-1}}{2}\frac{\partial^2}{\partial \epsilon_1^2}\right)\theta k_{1-}\cdot(\phi_1 + \Omega^{-1/2}\epsilon_1) \\ &\quad + \left(-\Omega^{-1/2}\frac{\partial}{\partial \epsilon_i} + \frac{\Omega^{-1}}{2}\frac{\partial^2}{\partial \epsilon_i^2}\right)k_2 + \left(\Omega^{-1/2}\frac{\partial}{\partial \epsilon_2} + \frac{\Omega^{-1}}{2}\frac{\partial^2}{\partial \epsilon_2^2}\right)k_{2-}(\phi_2 + \Omega^{-1/2}\epsilon_2) \\ &\quad + \left(-\Omega^{-1/2}\frac{\partial}{\partial \epsilon_i} + \frac{\Omega^{-1}}{2}\frac{\partial^2}{\partial \epsilon_i^2}\right)k_3(\phi_1 + \Omega^{-1/2}\epsilon_1)\cdot(\phi_2 + \Omega^{-1/2}\epsilon_2) \\ &\quad + \left(\Omega^{-1/2}\frac{\partial}{\partial \epsilon_3} + \frac{\Omega^{-1}}{2}\frac{\partial^2}{\partial \epsilon_3^2}\right)k_{3-}(\phi_3 + \Omega^{-1/2}\epsilon_3)\Bigg]\Pi(\epsilon_i,t).\end{aligned} \tag{13}$$

After ordering according to powers of $\Omega$, we get

$$\begin{aligned}&\frac{\partial \Pi(\epsilon_i,t)}{\partial t} - \sum_i \Omega^{1/2}\frac{\partial \Pi(\epsilon_i,t)}{\partial \epsilon_i}\frac{\partial \phi_i}{\partial t} \\ &= \Omega^{1/2}\left[(-\theta k_1 + \theta k_{1-}\phi_1)\frac{\partial \Pi}{\partial \epsilon_1} + (-k_2 + k_{2-}\phi_2)\frac{\partial \Pi}{\partial \epsilon_2} + (-k_3\phi 1\phi_2 + k_{3-}\phi_3)\frac{\partial \Pi}{\partial \epsilon_2}\right] + \\ &\Omega^0\Bigg[(\frac{k_1}{2} + \frac{\theta k_1\phi_1}{2})\frac{\partial^2 \Pi}{\partial \epsilon_1^2} + \theta k_1\epsilon_1\frac{\partial \Pi}{\partial \epsilon_1} + \theta k_1\Pi + (\frac{k_2}{2} + \frac{k_{2-}\phi_2}{2})\frac{\partial^2 \Pi}{\partial \epsilon_2^2} + k_{2-}\epsilon_2\frac{\partial \Pi}{\partial \epsilon_2} + k_{2-}\Pi \\ &+ (\frac{k_3\phi 1\phi_2}{2} + \frac{k_{3-}\phi_3}{2})\frac{\partial^2 \Pi}{\partial \epsilon_3^2} + (k_3\phi_2\epsilon_1 + k_3\phi_1\epsilon_2)\frac{\partial \Pi}{\partial \epsilon_3} + k_{3-}\epsilon_3\frac{\partial \Pi}{\partial \epsilon_3} + k_{3-}\Pi\Bigg] + \\ &O(\Omega^{-1/2}).\end{aligned} \tag{14}$$

Terms of order $\Omega^{1/2}$ contain the deterministic equations, are present on both sides and cancel. Hence, the remaining Fokker-Planck equation is

$$\begin{aligned}\frac{\partial \Pi(\epsilon_i,t)}{\partial t} =& (\frac{\theta k_1}{2} + \frac{\theta k_{1-}\phi_1}{2})\frac{\partial^2 \Pi}{\partial \epsilon_1^2} + \theta k_{1-}\epsilon_1\frac{\partial \Pi}{\partial \epsilon_1} + \theta k_1\Pi + (\frac{k_2}{2} + \frac{k_{2-}\phi_2}{2})\frac{\partial^2 \Pi}{\partial \epsilon_2^2} + k_{2-}\epsilon_2\frac{\partial \Pi}{\partial \epsilon_2} + k_{2-}\Pi \\ &+ (\frac{k_3\phi_1\phi_2}{2} + \frac{k_{3-}\phi_3}{2})\frac{\partial^2 \Pi}{\partial \epsilon_3^2} + (k_3\phi_2\epsilon_1 + k_3\phi_1\epsilon_2)\frac{\partial \Pi}{\partial \epsilon_3} + k_{3-}\epsilon_3\frac{\partial \Pi}{\partial \epsilon_3} + k_{3-}\Pi + O(\Omega^{-1/2}).\end{aligned} \tag{15}$$

From the Fokker-Planck equation we can calculate the second moments by multiplying equation 15 by the $\epsilon_i \epsilon_j$ and integrating over the whole domain

$$\frac{\partial}{\partial t}\int_{-\infty}^{\infty}\int_{-\infty}^{\infty}\epsilon_i\epsilon_j\Pi(\vec{\epsilon},t)d\epsilon =$$
$$\int_{-\infty}^{\infty}\int_{-\infty}^{\infty}\epsilon_i\epsilon_j\Bigg((\frac{\theta k_1}{2}+\frac{\theta k_{1-}\phi_1}{2})\frac{\partial^2\Pi}{\partial\epsilon_1^2}+\theta k_{1-}\epsilon_1\frac{\partial\Pi}{\partial\epsilon_1}+\theta k_1\Pi+(\frac{k_2}{2}+\frac{k_{2-}\phi_2}{2})\frac{\partial^2\Pi}{\partial\epsilon_2^2}$$
$$+k_{2-}\epsilon_2\frac{\partial\Pi}{\partial\epsilon_2}+k_{2-}\Pi+(\frac{k_3\phi_1\phi_2}{2}+\frac{k_{3-}\phi_3}{2})\frac{\partial^2\Pi}{\partial\epsilon_3^2}+(k_3\phi_2\epsilon_1+k_3\phi_1\epsilon_2)\frac{\partial\Pi}{\partial\epsilon_3}$$
$$+k_{3-}\epsilon_3\frac{\partial\Pi}{\partial\epsilon_3}+k_{3-}\Pi+O(\Omega^{-1/2})\Bigg)d\epsilon_i d\epsilon_j \quad (16)$$

to get the Matrix Differential Equation

$$\frac{\partial\mathbf{\Sigma}}{\partial t}=\mathbf{J\Sigma}+\mathbf{\Sigma J^T}+\mathbf{D}+\mathcal{O}(\mathbf{\Omega^{-3/2}}) \quad (17)$$

In steady state $\frac{\partial\mathbf{\Sigma}}{\partial t}=0$, equation 17 resembles a Lyapunov Equation, that can be solved with a computer algebra software (e.g. Mathematica comes with a built in function called LyapunovSolve). We are interested in the normalized covariance or noise $\frac{\langle\epsilon_i\epsilon_j\rangle}{\langle\phi_i\phi_j\rangle}$ of each species, thus we also normalize equation 17 by division with $\langle\phi_i\rangle\langle\phi_j\rangle$. For the noise in the output $i=j=3$ we get

$$\frac{\langle\epsilon_3\rangle^2}{\langle\phi_3\rangle^2}=\underbrace{\frac{1}{\langle\phi_3\rangle}}_{\text{inherent noise}}+\underbrace{\frac{1}{\langle\phi_1\rangle}\frac{k_{3-}}{(\theta k_{1-}+k_{3-})}}_{\text{transmitted from input}}+\underbrace{\frac{1}{\langle\phi_2\rangle}\frac{k_{3-}}{(k_{2-}+k_{3-})}}_{\text{transmitted from embedding}}. \quad (18)$$

The Fokker-Plank equation has the general form:

$$\frac{\partial\Pi}{\partial t}=-\sum_{i,w=1}J_{iw}\frac{\partial}{\partial\epsilon_i}(\epsilon_w\Pi)+\frac{1}{2}\sum_{i,k=1}D_{ik}\frac{\partial^2}{\partial\epsilon_i\partial\epsilon_k}\Pi+O(\Omega^{-1/2}) \quad (19)$$

with

$$J_{iw}=\sum_{j=1}^{R}S_{ij}\frac{\partial f_j(\vec{n},\Omega)}{\partial\phi_w} \qquad D_{ik\ldots r}=\sum_{j=1}^{R}S_{ij}S_{kj}\ldots S_{rj}f_j(\vec{n},\Omega). \quad (20)$$

For the studied system this gives the matrices

$$J_{iw}=\begin{pmatrix}\theta k_1 & 0 & 0\\ 0 & -k_{2-}E/\Omega & 0\\ k_3E/\Omega^2 & k_3I/\Omega^2 & -k_{3-}\end{pmatrix} \quad (21)$$

$$D_{ik\ldots r}=\begin{pmatrix}\theta k_1+\theta k_1 I & 0 & 0\\ 0 & k_2+k_{2-}E/\Omega & 0\\ 0 & 0 & k_3IE/\Omega^2+k_{3-}X/\Omega.\end{pmatrix} \quad (22)$$

Note, that while the mean levels of $I$ are unaffected by the input noise parameter $\theta$, both the Jacobian $J_{iw}$ and the Diffusionmatrix $D_{ik\ldots r}$ depend on it.

### Derivation of the autocorrelation

By multiplying both sides of 15 by $\epsilon(t)\epsilon(t+\tau)$ and integrating over the whole domain we have

$$\frac{\partial\langle\epsilon_i(t)\epsilon_j(t+\tau)\rangle}{\partial t} = \int_{-\infty}^{\infty}\int_{-\infty}^{\infty}\epsilon_i(t)\epsilon_j(t+\tau)\frac{\partial\Pi(\epsilon_i,t)}{\partial t}d\epsilon_i d\epsilon_j$$

$$= \int_{-\infty}^{\infty}\int_{-\infty}^{\infty}\epsilon_i(t)\epsilon_j(t+\tau)\Bigg[(\frac{\theta k_1}{2}+\frac{\theta k_{1-}\phi_1}{2})\frac{\partial^2\Pi}{\partial\epsilon_1^2}+\theta k_{1-}\epsilon_1\frac{\partial\Pi}{\partial\epsilon_1}$$
$$+\theta k_{1-}\Pi+(\frac{k_2}{2}+\frac{k_{2-}\phi_2}{2})\frac{\partial^2\Pi}{\partial\epsilon_2^2}+k_{2-}\epsilon_2\frac{\partial\Pi}{\partial\epsilon_2}+k_{2-}\Pi$$
$$+(\frac{k_3\phi 1\phi_2}{2}+\frac{k_{3-}\phi_3}{2})\frac{\partial^2\Pi}{\partial\epsilon_3^2}+(k_3\phi_2\epsilon_1+k_3\phi_1\epsilon_2)\frac{\partial\Pi}{\partial\epsilon_3}$$
$$+k_{3-}\epsilon_3\frac{\partial\Pi}{\partial\epsilon_3}+k_{3-}\Pi+O(\Omega^{-1/2})\Bigg]d\epsilon_i d\epsilon_j. \quad (23)$$

For $i=j$ the above integral gives the autocorrelation, for $i=1$ we have

$$\frac{\partial\langle\epsilon_1(t)\epsilon_1(t+\tau)\rangle}{\partial t} = \int_{-\infty}^{\infty}\int_{-\infty}^{\infty}\epsilon_1(t)\epsilon_1(t+\tau)\left[(\frac{\theta k_1}{2}+\frac{\theta k_{1-}\phi_1}{2})\frac{\partial^2\Pi}{\partial\epsilon_1^2}+\theta k_{1-}\epsilon_1\frac{\partial\Pi}{\partial\epsilon_1}+\theta k_{1-}\Pi\right]d\epsilon_1 d\epsilon_1. \quad (24)$$

Only the drift terms (first derivatives $\frac{\partial\Pi}{\partial\epsilon_1}$) contribute and using $dt=d\tau$ we have

$$\frac{\partial\langle\epsilon_1(t)\epsilon_1(t+\tau)\rangle}{\partial\tau} = -2\theta k_{1-}\langle\epsilon_1(t)\epsilon_1(t+\tau)\rangle+\theta k_{1-}\langle\epsilon_1(t)\epsilon_1(t+\tau)\rangle \quad (25)$$

with the exponential solution

$$\langle\epsilon_1(t)\epsilon_1(t+\tau)\rangle = \langle\epsilon_1(t)\epsilon_1(t)\rangle e^{-\theta k_{1-}\tau} \quad (26)$$

where the initial condition is determined by the covariance $\langle\epsilon_1(t)\epsilon_1(t)\rangle$. Similarly for $\epsilon_2$ we get

$$\langle\epsilon_2(t)\epsilon_2(t+\tau)\rangle = \langle\epsilon_2(t)\epsilon_2(t)\rangle e^{-k_{2-}\tau}. \quad (27)$$

From the Master Equation we get for $\langle\epsilon_3(t)\epsilon_3(t+\tau)\rangle$

$$\frac{\partial\langle\epsilon_3(t)\epsilon_3(t+\tau)\rangle}{\partial t} = \int_{-\infty}^{\infty}\int_{-\infty}^{\infty}\epsilon_3(t)\epsilon_3(t+\tau)\Bigg[(\frac{k_3\phi 1\phi_2}{2}+\frac{k_{3-}\phi_3}{2})\frac{\partial^2\Pi}{\partial\epsilon_3^2}$$
$$+(k_3\phi_2\epsilon_1+k_3\phi_1\epsilon_2)\frac{\partial\Pi}{\partial\epsilon_3}+k_{3-}\epsilon_3\frac{\partial\Pi}{\partial\epsilon_3}+k_{3-}\Pi\Bigg]d\epsilon_3 d\epsilon_3 \quad (28)$$

this drift term also contains the interactions so integration gives

$$\frac{\partial\langle\epsilon_3(t)\epsilon_3(t+\tau)\rangle}{\partial\tau} = 0-k_3\phi_2\langle\epsilon_1(t)\epsilon_3(t+\tau)\rangle-k_3\phi_1\langle\epsilon_2(t)\epsilon_3(t+\tau)\rangle$$
$$-2k_{3-}\langle\epsilon_3(t)\epsilon_3(t+\tau)\rangle+k_{3-}\langle\epsilon_3(t)\epsilon_3(t+\tau)\rangle \quad (29)$$

which depends on $\langle\epsilon_1(t)\epsilon_3(t+\tau)\rangle$ and $\langle\epsilon_2(t)\epsilon_3(t+\tau)\rangle$. Similarly integration of the master equation

$$\frac{\partial\langle\epsilon_1(t)\epsilon_3(t+\tau)\rangle}{\partial t} = \int_{-\infty}^{\infty}\int_{-\infty}^{\infty}\epsilon_1(t)\epsilon_3(t+\tau)\Bigg[(\frac{\theta k_1}{2}+\frac{\theta k_{1-}\phi_1}{2})\frac{\partial^2\Pi}{\partial\epsilon_1^2}+\theta k_{1-}\epsilon_1\frac{\partial\Pi}{\partial\epsilon_1}+\theta k_{1-}\Pi$$
$$+(\frac{k_3\phi 1\phi_2}{2}+\frac{k_{3-}\phi_3}{2})\frac{\partial^2\Pi}{\partial\epsilon_3^2}+(k_3\phi_2\epsilon_1+k_3\phi_1\epsilon_2)\frac{\partial\Pi}{\partial\epsilon_3}+k_{3-}\epsilon_3\frac{\partial\Pi}{\partial\epsilon_3}+k_{3-}\Pi\Bigg]d\epsilon_3 d\epsilon_3 \quad (30)$$

yields

$$\frac{\partial\langle\epsilon_1(t)\epsilon_3(t+\tau)\rangle}{\partial\tau} = -2\theta k_{1-}\langle\epsilon_1(t)\epsilon_3(t+\tau)\rangle + \theta k_{1-}\langle\epsilon_1(t)\epsilon_3(t+\tau)\rangle + 0 \tag{31}$$

and solving the ordinary differential equation gives

$$\langle\epsilon_1(t)\epsilon_3(t+\tau)\rangle = \langle\epsilon_1(t)\epsilon_3(t)\rangle e^{-\theta k_{1-}\tau}. \tag{32}$$

The same procedure for the other interaction results in

$$\langle\epsilon_2(t)\epsilon_3(t+\tau)\rangle = \langle\epsilon_2(t)\epsilon_3(t)\rangle e^{-k_{2-}\tau}. \tag{33}$$

The complementary solution to the ordinary differential equation 28 for $\langle\epsilon_3(t)\epsilon_3(t+\tau)\rangle$ without interactions

$$\frac{\partial\langle\epsilon_3(t)\epsilon_3(t+\tau)\rangle}{\partial\tau} + k_{3-}\langle\epsilon_3(t)\epsilon_3(t+\tau)\rangle = 0 \tag{34}$$

is simply

$$\langle\epsilon_3(t)\epsilon_3(t+\tau)\rangle_c = Ce^{-k_{3-}\tau}. \tag{35}$$

Considering inhomogeneity from interactions we have

$$\frac{\partial\langle\epsilon_3(t)\epsilon_3(t+\tau)\rangle}{\partial\tau} + k_{3-}\langle\epsilon_3(t)\epsilon_3(t+\tau)\rangle = -k_3\phi_2\langle\epsilon_1(t)\epsilon_3(t+\tau)\rangle - k_3\phi_1\langle\epsilon_2(t)\epsilon_3(t+\tau)\rangle \tag{36}$$

and the particular solution is

$$\langle\epsilon_3(t)\epsilon_3(t+\tau)\rangle_p = e^{-k_{3-}\tau}\int(-\phi_2\langle\epsilon_1(t)\epsilon_3(t)\rangle e^{-\theta k_{1-}\tau} - \phi_1\langle\epsilon_2(t)\epsilon_3(t)\rangle e^{-k_{2-}\tau})e^{k_{3-}\tau}d\tau \tag{37}$$

$$= \frac{\phi_2}{(k_{3-}-\theta k_{1-})}\langle\epsilon_1(t)\epsilon_3(t)\rangle e^{-\theta k_{1-}\tau} + \frac{\phi_1}{(k_{3-}-k_{2-})}\langle\epsilon_2(t)\epsilon_3(t)\rangle e^{-k_{2-}\tau}. \tag{38}$$

Normalizing and inserting the covariances, we get

$$\frac{\langle\epsilon_3(t)\epsilon_3(t+\tau)\rangle_p}{\langle\phi_3\rangle^2} = \frac{1}{\phi_1}\frac{k_{3-}}{(\theta k_{1-}+k_{3-})}\frac{\phi_2}{(k_{3-}-\theta k_{1-})}e^{-\theta k_{1-}\tau} + \frac{1}{\phi_2}\frac{k_{3-}}{(k_{2-}+k_{3-})}\frac{\phi_1}{(k_{3-}-k_{2-})}e^{-k_{2-}\tau} \tag{39}$$

And the general solution is

$$\frac{\langle\epsilon_3(t)\epsilon_3(t+\tau)\rangle}{\langle\phi_3\rangle^2} = \left(\frac{1}{\phi_3} + \frac{1}{\phi_1}\frac{k_{3-}}{(\theta k_{1-}+k_{3-})}\left(1-\frac{\phi_2}{(k_{3-}-\theta k_{1-})}\right) - \frac{1}{\phi_2}\frac{k_{3-}}{(k_{2-}+k_{3-})}\left(1-\frac{\phi_1}{(k_{3-}-k_{2-})}\right)\right)e^{-k_{3-}\tau}, \tag{40}$$

where we have chosen the integration constant $C$ such that for $\tau = 0$, we recover the covariance. Rearranging using $A = \frac{\phi_2}{(k_{3-}-\theta k_{1-})}$ and $B = \frac{\phi_1}{(k_{3-}-k_{2-})}$ yields

$$\frac{\langle\epsilon_3(t)\epsilon_3(t+\tau)\rangle_{tot}}{\langle\phi_3\rangle^2} = \underbrace{\frac{1}{\phi_3}e^{-k_{3-}\tau}}_{\text{inherent}} + \underbrace{\frac{1}{\phi_1}\frac{k_{3-}}{(\theta k_{1-}+k_{3-})}(Ae^{-\theta k_{1-}\tau} + (1-A)e^{-k_{3-}\tau})}_{\text{from input}} \tag{41}$$

$$+ \underbrace{\frac{1}{\phi_2}\frac{k_{3-}}{(k_{2-}+k_{3-})}(Be^{-k_{2-}\tau} + (1-B)e^{-k_{3-}\tau})}_{\text{embedding}}. \tag{42}$$

### Gillespie Simulations of the Reaction System

To compare the approximate results obtained by the LNA, Gillespie simulations were performed. At least $10^3$ simulations with $10^5$ iterations were run for 8 different input noise parameter values $\theta$ and 8 different input mean levels (Figure S1). The steady state mean and variability of each species was extracted from the time course data of each simulation and the mean noise $(CV^2)$ plotted against the parameter values. The variability between extracted variabilities of each of the $10^3$ simulation was used to estimate the simulation error seen in Figure 3 and 4.

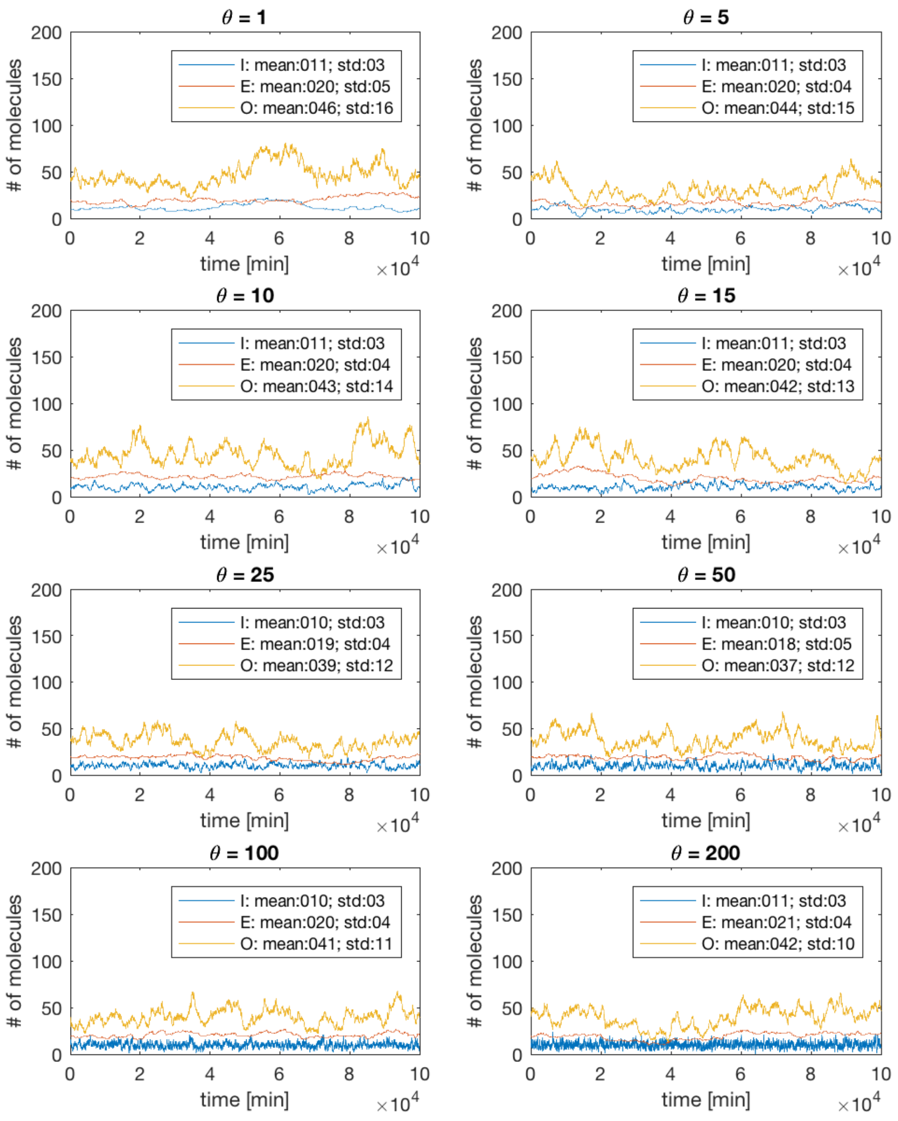


Figure S1: Gillespie simulation time courses for 1 and parameters: $\Omega = 1, k_1 = 10^{-5}, k_{1-} = 10^{-6}, k_2 = 2 * 10^{-5}, k_{2-} = 10^{-6}, k_3 = 2 * 10^{-6}, k_{3-} = 10^{-5}$. Each plot shows how the molecule numbers evolve over time for one set of parameter values. The change of time scale in the input trajectory (blue) from one parameter set to the other is clearly visible.

### Fitting Hill Functions to Time Course Data

After tracking HeLa cell nuclei, nuclear Höchst dye intensities were extracted and plotted over time (see Figure S2). Sudden increase of the signal marks cell death. Initial tries to identify the time of death by determining the time, when the Höchst intensity crossed a simple threshold turned out not to be robust, because the absolute range of intensities measured across time varied a lot between cells. In other words some cells start above the threshold, while others ended below. The intensity jumps during death though, are very robust.

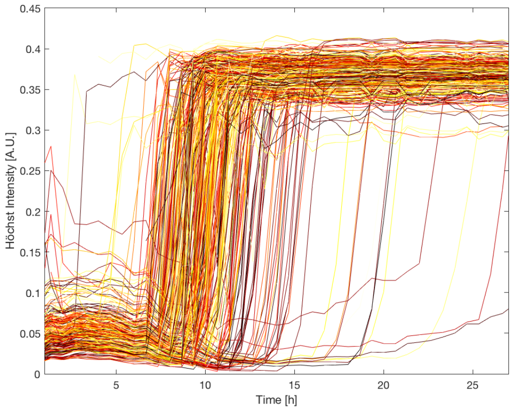


Figure S2: The horizontal axis represents time after cisplatin treatment and the vertical axis the intensity of the Höchst staining in arbitrary units. The plot shows time courses of Höchst dye intensities for hundreds of single cells under cisplatin treatment. Sudden signal increases indicate cell death.

To improve the robustness of death time detection, Hill functions were fitted to the time course data (see Figure S3). Parameters where fitted with the inbuilt Matlab nonlinear fit function (nlinfit) and the model $@(b, x)(b(3) + b(1) * x.^{15}./(b(2)^{15} + x^{15}))$. This function allows to tune the minimum $b(3)$ and maximum $b(1)$ of the signal increase and the time when the signal increase occurs $b(2)$. The latter $b(2)$ parameter fits can be reliably used for fully automated detection of death times.

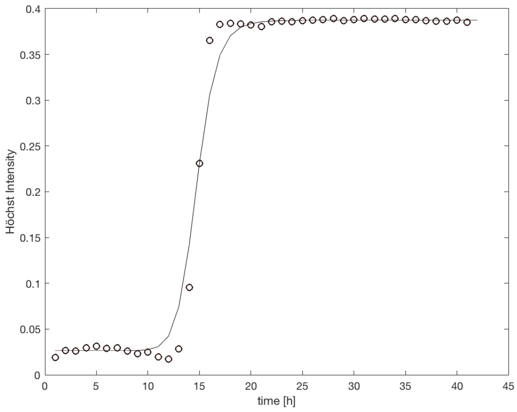


Figure S3: The horizontal axis represents time after cisplatin treatment and the vertical axis the intensity of the Höchst staining in arbitrary units. The plot shows an example of a fit of a Hill Function (continuous curve) to the Höchst time course data (circles) of a single cell.

[2] Edda Klipp, Wolfram Liebermeister, Christoph Wierling, Axel Kowald, Hans Lehrach, and Ralf Herwig. Systems Biology: A Textbook. Wiley-VCH, 2009.
